# The Role of Human-Specific lncRNA in Hyaline Cartilage Development

**DOI:** 10.64898/2026.02.17.706478

**Authors:** Tatsunori Osone, Tomoka Takao, Takeshi Takarada

## Abstract

One of the distinctive characteristics of humans is their bipedalism. To achieve upright bipedal walking, the angles of the pelvis and femur have been altered. Although evolutionary hypotheses on the transition to bipedalism exist, the molecular mechanisms remain unclear. This study attempts to elucidate these mechanisms using a system for inducing hyaline cartilage-like tissue from human iPS cells via limb bud like mesenchymal cells. Focus was placed on non-coding RNAs, known for their potential in generating biological diversity. Bulk RNA sequencing was conducted to compare the expression and functions of human-specific long non-coding RNAs between limb bud like mesenchymal cells and induced hyaline cartilage-like tissue. The results indicated that human-specific lncRNAs, significantly upregulated in hyaline cartilage-like tissue, may regulate genes related to the extracellular matrix. These findings suggest the potential to develop regenerative cartilage tissue with enhanced ECM quality through controlling human-specific lncRNAs. Additionally, studying human-specific lncRNAs could elucidate mechanisms of diseases that are less common in other species but more prevalent in humans.

## Introduction

Humans possess a markedly different body structure compared to other organisms. The human brain is notably larger than that of other animals, with a particularly well-developed frontal lobe(Hofman, 2014). Humans possess numerous facial muscles, enabling rich expression of emotions(Burrows, 2008). The throat and vocal cords can produce complex sounds, facilitating language-based communication(Nishimura et al., 2022). The hands exhibit opposable thumbs, permitting intricate manipulation(Richmond et al., 2016). Humans have less body hair and more developed sweat glands(Kamberov et al., 2018). Additionally, humans are the only species that practises upright bipedalism.

In evolutionary biology, several hypotheses have been proposed to explain the attainment of upright bipedalism. These include the Savannah Hypothesis, the Carrying Hypothesis, and the Thermoregulatory Hypothesis(Harcourt-Smith, 2010). The Savannah Hypothesis attributes bipedalism to environmental changes. Approximately six million years ago, the African forests contracted, giving rise to expansive savannahs. Consequently, human ancestors needed to traverse open grasslands, and it is believed that upright bipedalism allowed them to efficiently survey large areas. The Carrying Hypothesis suggests that the freeing of the hands enabled the transport of food, tools, and offspring, which was advantageous for survival and reproduction. The use of tools is considered a factor in human evolution, contributing to the development of intelligence. The Thermoregulatory Hypothesis posits that bipedalism was an adaptation for survival in hot environments. By standing upright, the body reduced the surface area exposed to direct solar radiation and heat reflected from the ground. Moreover, the increased surface area exposed to wind facilitated effective heat dissipation.

Despite the various hypotheses proposed in evolutionary biology, the molecular biological mechanisms underlying human bipedalism remain elusive. The evolution of bipedalism in humans is closely related to changes in the structure of the lower limbs. It is well known that the lower limbs of humans are significantly longer compared to those of the Pan genus, and there are notable differences in the angle of the pelvis(O’Neill et al., 2015). Therefore, elucidating the human-specific mechanisms during the developmental process when the skeleton is formed could lead to the formulation of hypotheses regarding the molecular biological mechanisms of evolution. The skeleton is formed through endochondral ossification, where cartilage is replaced by bone, and secondary ossification occurs at the epiphyses(Long and Ornitz, 2013). Hence, it is essential to focus on human-specific cartilage formation during development. Previously, a method was established for differentiating human iPS cells into expandable limb bud like mesenchymal cells (ExpLBM) via lateral plate mesoderm, and subsequently into hyaline cartilage-like tissue (HCT)(Yamada et al., 2021). Since this method mimics the developmental process, it was considered that comparing RNA expression between ExpLBM and HCT could reveal RNAs involved in human-specific cartilage formation.

However, since mRNA exhibits a high degree of sequence conservation across species, it might be challenging to elucidate the molecular biological mechanisms using mRNA alone. On the other hand, long non-coding RNAs (lncRNAs), which are RNA molecules longer than 200 nucleotides that do not encode proteins, show less sequence conservation compared to mRNA(Makałowski et al., 1996; Noviello et al., 2018). Moreover, lncRNAs are known to exhibit more tissue-specific expression patterns than mRNA, suggesting significant roles in cell type-specific processes(Statello et al., 2021). Unlike the approximately 20,000 types of mRNA, the number of lncRNAs reaches hundreds of thousands, and due to alternative splicing, the number of isoforms increases dramatically(“RNAcentral: a comprehensive database of non-coding RNA sequences,” 2017). This complexity makes functional evaluation challenging, and consequently, lncRNA research is not as advanced. lncRNAs play crucial roles in various cellular processes, such as the cell cycle, differentiation, and metabolism. Their mechanisms involve interactions with DNA, RNA, and proteins, leading to the regulation of epigenetic modifications, transcription, translation, post-translational modifications, and the stability of proteins and RNAs(Herman et al., 2022). Particularly in transcriptional regulation, lncRNAs are known to cis-acting by regulating the expression of genes located near their own genetic loci, or by binding to promoter and enhancer regions of target genes to control transcription(Gil and Ulitsky, 2020). This regulation occurs by either inhibiting or promoting the binding of transcription factors and the transcription initiation complex.

LncRNAs are known to interact with various biomolecules; consequently, numerous tools have been developed to predict their functions. For predicting interactions with DNA, tools such as Triplex Domain Finder (TDF) and PATO have been developed(Amatria-Barral et al., 2023; Kuo et al., 2019). TDF utilises an efficient bit-parallel algorithm and a concept akin to motif overrepresentation analysis to rapidly predict regions where lncRNAs form triple helices with DNA. For predicting interactions with proteins and RNA, tools such as linc2function and EDLMFC have been developed(Ramakrishnaiah et al., 2023; Wang et al., 2021). Linc2function is a tool that extracts features through machine learning from nucleotide sequences and predicts functional characteristics via interactome and secondary structure analysis. It can be executed locally or through a web server.

By utilising the characteristics of lncRNAs and these tools, it is hypothesised that the relationship between human-specific lncRNAs and the evolution of bipedalism, which is a significant difference between humans and other primates such as Pan, can be elucidated. Therefore, in this study, an attempt was made to evaluate the functions of cartilage-related lncRNAs from an evolutionary perspective by predicting the roles of human-specific lncRNAs that are significantly up-regulated in chondrocytes.

## Materials and methods

### Cell culture and Differentiation

The methods employed were based on those outlined in our previous publication(Yamada et al., 2021). Briefly, human induced pluripotent stem cells (414C2 PRRX1-tdTomato; 3 × 10^4^ cells) were suspended in 1 ml of StemFit (Ajinomoto) containing 10 μM Y27632 (Wako) and added to a 3.5 cm culture dish containing 4 μl iMatrix511. The culture medium was replaced the next day with fresh StemFit without Y27632. After culturing for 2 d, the cells were washed with PBS and differentiation was induced by changing the culture medium at each time point. Accutase (Thermo Fisher) was used to dissociate LBM-like cells, and 2 × 10^5^ cells were suspended in ExpLBM medium containing 16 μl iMatrix511 and then added to a 6 cm culture dish. The culture media were replaced with fresh ExpLBM medium every 2 d. To induce chondrogenic induction using pellet culture conditions (3DCI), 1 × 10^5^ ExpLBM cells were suspended in 200 μl chondrogenic culture step 1 medium and added to 96-well ultralow U-bottom plates (Corning). The cells were pelleted by centrifugation at 2,000 r.p.m. for 5 min. At the end of steps 1 and 2, the culture media were changed to chondrogenic culture step 2 and step 3 media, respectively.

### Bulk RNA-Seq

Total RNA was extracted using an RNeasy kit (Qiagen), and sequencing libraries were prepared using a KAPA RNA HyperPrep Kit with RiboErase (HMR) (Kapa Biosystems, USA) and a SeqCap Adapter Kit (Set A or Set B, Roche, USA) according to the manufacturer’s instructions. Sequencing libraries were transferred to AZENTA (Suzhou, China) and were loaded onto a HiSeq 2500 system (Illumina, USA) for sequencing. All sequence reads were extracted in FASTQ format using the CASAVA 1.8.4 pipeline. Fastp (version 0.23.4) was used to remove adapters and filter raw reads of < 60 bases in addition to leading and trailing bases with a read quality of less than 30. For the pseudoalignment of the filtered reads to the GRCh38.p14, Kallisto (version 0.48.0) was used. GeTMM normalization was performed using edgeR (version 4.0.3) to account for sample variation. Differentially expressed genes were identified through NOISeq (version 2.46.0) analysis. The raw RNA-seq data were deposited in the NCBI GEO database under accession number GSE165620, .

### Databases

RNACentral(“RNAcentral: a comprehensive database of non-coding RNA sequences,” 2017) was utilised for the identification of lncRNAs. For the identification of human-specific lncRNAs, LncBook2.0(Li et al., 2023) was employed.

### Gene Ontology Analysis

Gene Ontology (GO) analysis was performed using the clusterProfiler package (v4.10.1) in the R (v4.3.3) environment against the org.Hs.eg.db database (v3.18.0). The parameters used for the analysis were ont=“ALL”, pAdjustMethod=“BH”, pvalueCutoff=0.05, and qvalueCutoff=0.0. Visualisation was conducted using the ggplot2 package (v3.5.0).

### lncRNA Function Prediction

To predict the proteins binding to lncRNAs, the linc2function web server(Ramakrishnaiah et al., 2023) was utilised with parameters set to ANN, HumanSpecific, and Full Model. For peptide function prediction, DeepGOWeb (v1.0.18)(Kulmanov et al., 2021) was used, with the prediction threshold set to the default value of 0.3. Triplex prediction was conducted using the TDF(Kuo et al., 2019) included in the Regulatory Genomics Toolbox (v1.0.2), executed in promotertest mode against the hg38 genome assembly. All parameters were set to their default values.

## Results

### lncRNA Profiles During Chondrogenic Differentiation

Bulk RNA-Seq was performed on ExpLBM and HCT (Fig. 1A). From the obtained RNA expression profiles, non-coding RNAs were extracted, and MA plots were generated (Fig. 1B). There were 1,346 HCT-enriched lncRNAs (log2FC ≥ 1, prob > 0.8) and 1,082 ExpLBM-enriched lncRNAs (log2FC ≤ -1, prob > 0.8). Known cartilage-related lncRNAs were significantly highly expressed in HCT (Table 1). There were no significant differences in the expression of other lncRNAs reported in the reference literature. Although the referenced paper also reports on microRNAs, small RNA-Seq was not conducted; hence, the quantification was based on the expression levels of pri-miRNA and miRNA host gene. To extract human-specific lncRNAs from the HCT-enriched lncRNAs, the LncBook2.0 database, which includes conservation information, was used (Fig. 1C). As a result, 740 lncRNAs were identified (Fig. 1D, Supp. Fig. 1). Interestingly, the greatest number of conserved lncRNAs was found among the Eutheria, while those conserved for a longer period, excluding those conserved among the Euteleostomi, were present in similar quantities. Additionally, the lncRNAs conserved for a longer period were predominantly known to encode peptides, as identified by Ribo-Seq, whereas the more recently evolved lncRNAs post-Hominoidea scarcely encoded peptides. Furthermore, despite the relatively lower quantity of lncRNAs conserved between Primates and Hominini, a higher number of Homo-specific lncRNAs were observed in comparison.

**Figure 1.**
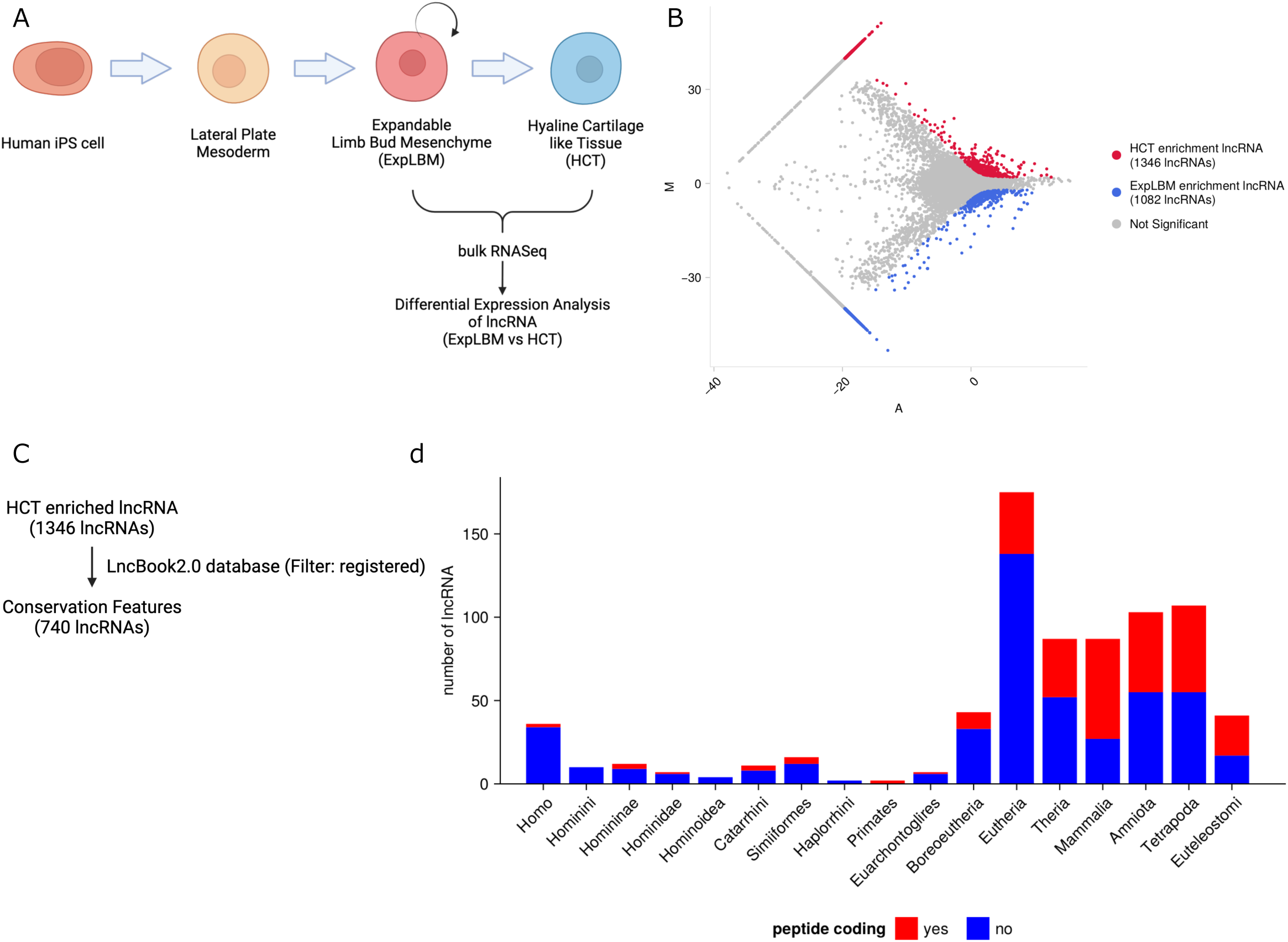
lncRNA Expression Profiles. (A) Schematic overview of the methodology used in this study. (B) MA plots comparing ExpLBM and HCT respective lncRNA expression levels. red: HCT enrichment ncRNA, blue: ExpLBM enrichment ncRNA. (C) Methodology for obtaining the conservation features. (D) Bar plot of Conservation features of DEL_HCT. The number of human lncRNAs per phylogenetic category is plotted against 740 HCT-enriched lncRNAs, classified by conservation age on the horizontal axis and by phylogenetic category on the vertical axis.

**Table 1.**
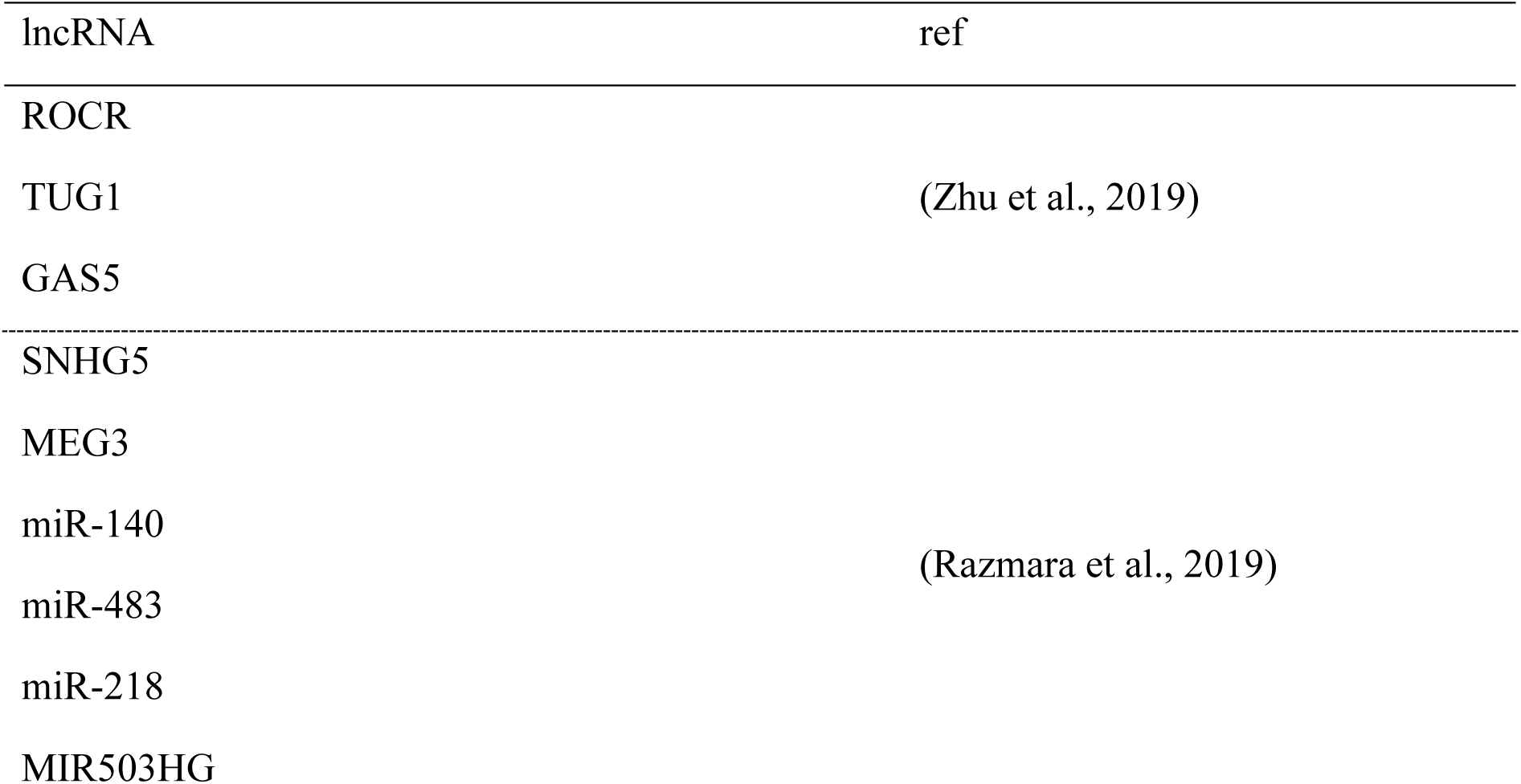
known cartilage-associated lncRNAs.

### Functional Prediction of Human-Specific lncRNAs

Among the differentially expressed lncRNAs (DELs) between ExpLBM and HCT, 36 human-specific lncRNAs (DEL_HCT_hs) were identified as having significantly increased expression levels in HCT. Among DEL_HCT_hs, HSALNG0113101 and HSALNG0142302 are known to be translated into peptides, capable of translating into 5 and 10 types of peptides, respectively (Suppl. Fig. 2A, 2B). Functional predictions for these 15 peptides were conducted using DeepGOWeb (Suppl. Fig. 2C). Only two peptides derived from HSALNG0142302 could have their functions predicted. Both were associated with the same Gene Ontology Molecular Function (GOMF) and Biological Process (GOBP), with GOMF related to binding functions and GOBP related to responses to stress and stimuli.

Additionally, the prediction of proteins binding to DEL_HCT_hs was conducted using linc2function for all proteins. As a result, it was predicted that 28 proteins would bind to 14 DEL_HCT_hs (Fig. 2A). EIF4B, FUS, MBNL1, and SFR1 were predicted to have binding potential with all the identified lncRNAs, whereas some proteins, such as A2BP1, were predicted to bind only to specific lncRNAs (Fig. 2B). Gene Ontology analysis of the 28 proteins predicted to bind to the 14 DEL_HCT_hs indicated that they are involved in RNA splicing and mRNA metabolic processes (Fig. 2C). Among the top 10 terms by GeneRatio, six functions involved in interactions with mRNA were identified: RNA splicing, regulation of mRNA metabolic process, RNA transport, establishment of RNA localization, mRNA stabilization, and positive regulation of translation.

**Figure 2.**
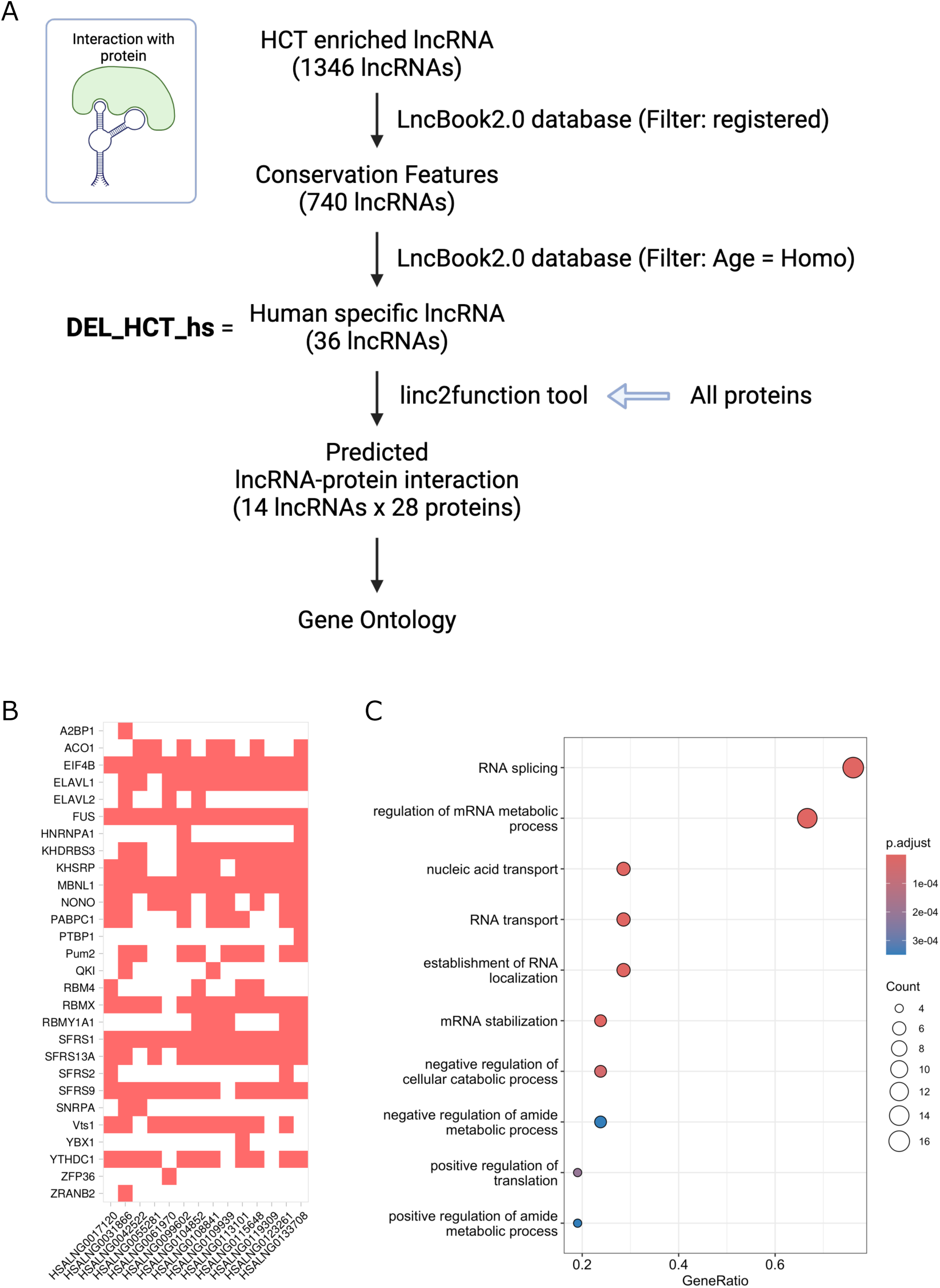
DEL_HCT_hs-Protein Interactions. (A) Schematic overview of the prediction methodology. (B) Correspondence map of predicted binding proteins with lncRNAs. Red indicates predicted binding pairs; white indicates non-predicted binding pairs. (C) Bubble plot of GO analysis results for 28 proteins predicted to bind lncRNAs

### Prediction of Gene Expression Regulation by Human-Specific lncRNA

To investigate whether DEL_HCT_hs could regulate the expression of the 2,841 mRNAs (DEG_HCT), which were significantly upregulated in HCT compared to ExpLBM, the interactions between the promoter regions of DEG_HCT and DEL_HCT_hs were predicted using TDF (Fig. 3A). The results indicated that 9 DEL_HCT_hs could form triplex structures with the promoter regions of DEG_HCT (Table 2). Among these, HSALNG0055279 was found near SMOC2, and HSALNG0123261 was located near the transcription factor NFIC. Using the NFIC GeneSet from Harmonizome 3.0, target genes of NFIC were identified, including the chondrocyte marker genes COL2A1 and SOX9, as well as the joint marker genes BARX1, COL14A1, EPS8, GEM, and NFIA. Analysis of the number of target genes for each lncRNA revealed that lnc-GOLGA6L4-2_4-HSALNG0107783_2 had the most targets, while HSALNG0123261 had the fewest (Fig. 3B). It should be noted that lncRNA names connected by a hyphen, such as lnc-GOLGA6L4-2_4-HSALNG0107783_2, represent different names from various databases, with the underscore being used to distinguish variants. Additionally, HSALNG0055648 had the most total triple helix-forming regions, and HSALNG0061970 had the fewest. Examination of genes with many triple helix-forming regions with DEL_HCT_hs revealed a high number of such regions between HSALNG0151253 and COL6A1 (Suppl. Fig. 3A). Focusing on chondrocyte marker genes and joint marker genes, it was predicted that ACAN, COL2A1, COL9A1, COL11A2, COMP, and SOX5 among the chondrocyte marker genes, and ABI3BP, BARX1, CHI3L1, EPS8, GEM, NFIA, THBS4, and TMEM30B among the joint marker genes, would form triplexes. While some genes, such as COL2A1, BARX1, EPS8, and THBS4, had very few triplex formation regions compared to other genes, there were also genes like CHI3L1 and TMEM30B that were predicted to form triplexes with all nine types (Fig. 3C, 3D). The DEL_HCT_hs with the highest number of triplex formation regions with chondrocyte marker genes was lnc-GOLGA6L4-2, whereas the DEL_HCT_hs with the highest number of triplex formation regions with joint marker genes were HSALNG0055648 and HSALNG0108834. Among the human-specific proteins within DEG_HCT, it was shown that GSTT2B, ZFP36L1, and ANGPTL5 could form triple helices with DEL_HCT_hs (Suppl. Fig. 3B). Among them, ANGPTL5 exhibited the highest number of total triplex formation regions with HSALNG0055648 and HSALNG0108834. Information gathered from GeneCards indicated that GSTT2B catalyses the conjugation of reduced glutathione to various electrophilic and hydrophobic compounds, ZFP36L1 functions as a transcription factor, and ANGPTL5 is predicted to act in the extracellular matrix and extracellular space containing collagen. Despite attempts to identify the target genes of ZFP36L1 using JASPAR, ZFP36L1 was not registered in the JASPAR database.

**Figure 3.**
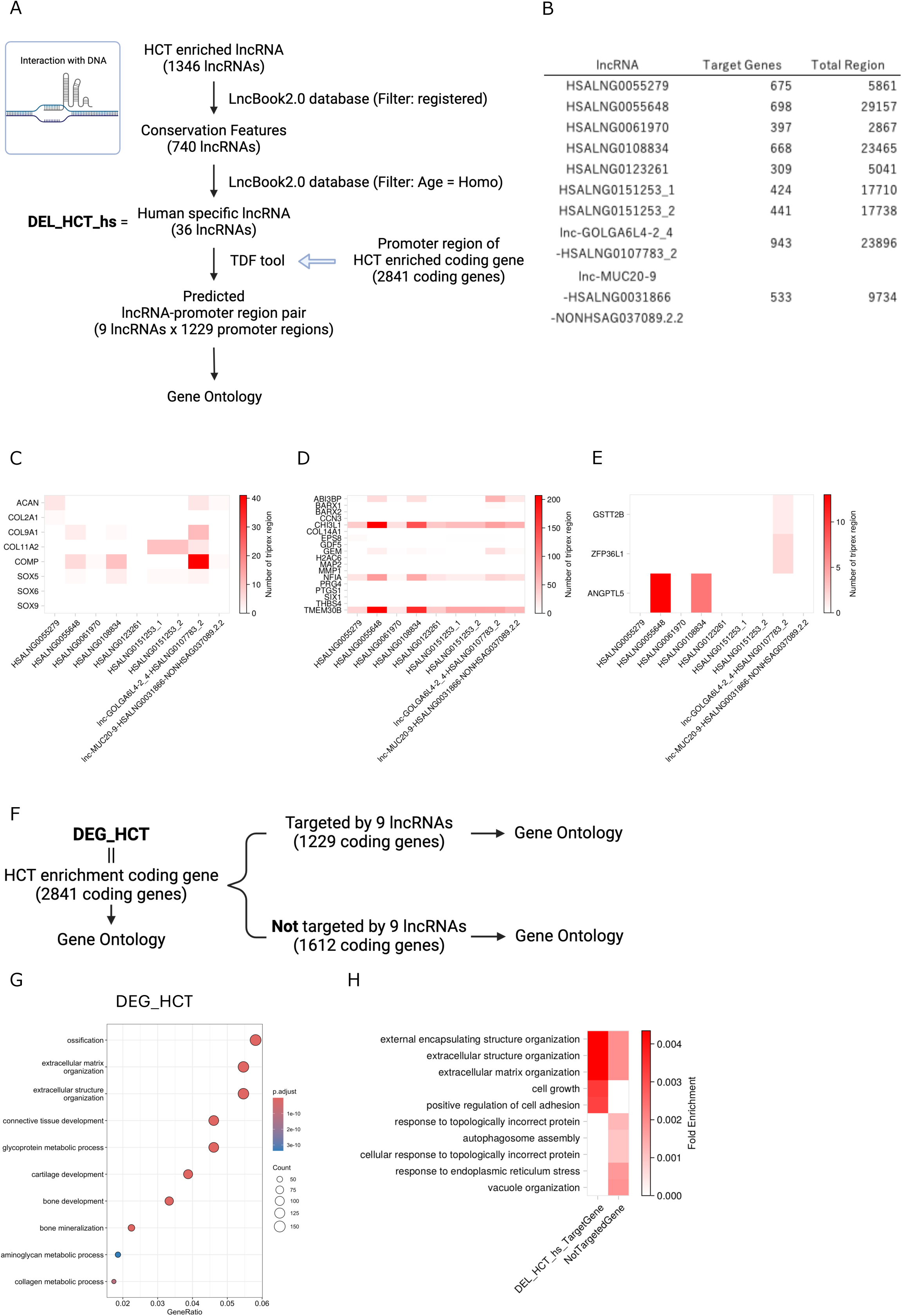
Triplex Predictions of DEL_HCT_hs. (A) Schematic overview of the prediction methodology. (B) Predicted lncRNAs forming triplex structures with DEG_HCT promoter regions, their target gene counts, and total formation sites. (C) Heatmap of formation sites in cartilage marker genes, plotted using the raw values of region numbers instead of Z-scores. (D) Heatmap of formation sites in chondrocyte marker genes, plotted using the raw values of region numbers instead of Z-scores. (E) Heatmap of formation sites in human-specific coding genes, plotted using the raw values of region numbers instead of Z-scores. (F) Subjects of Gene Ontology analysis. (G) Bubble plots representing the results of GO analysis in DEG_HCT. (H) The heatmap depicting Fold Enrichment for the GO analysis conducted on DEG_HCT predicted to be targeted by DEL_HCT_hs at the promoter region and other DEG_HCT. The top five and bottom five values of the log2(Fold Change of Fold Enrichment) were extracted, respectively.

**Table 2.**
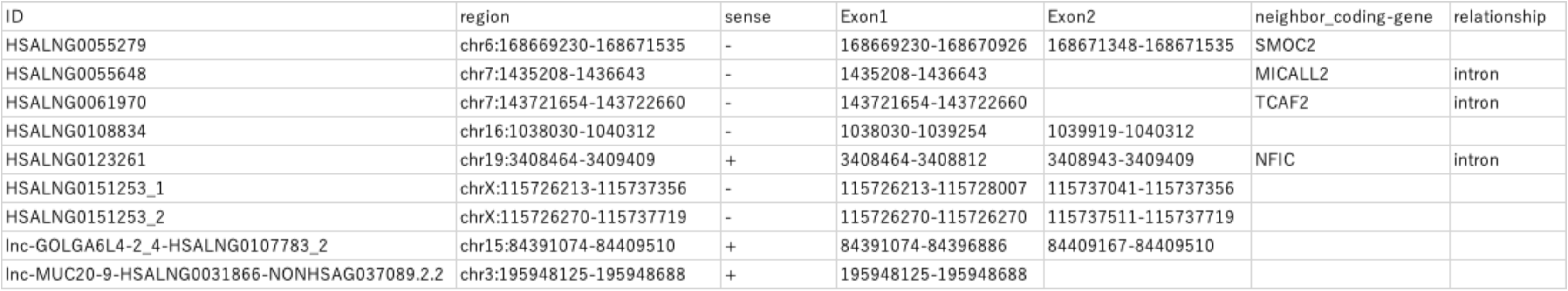
xxxxx

Given the potential of DEL_HCT_hs to form triple helices in the promoter regions of genes associated with cartilage, joints, and ECM, GO analysis was performed on DEG_HCT, the 1,229 genes predicted to form triple helices with DEL_HCT_hs in their promoter regions, and the remaining 1,612 genes not predicted to form triple helices with any DEL_HCT_hs (Fig. 3F). In DEG_HCT, ontologies related to ECM and bone/cartilage were enriched (Fig. 3G). To investigate the characteristics of the genes targeted by DEL_HCT_hs, a comparison was made between the genes predicted to form triple helices with DEL_HCT_hs in their promoter regions and the genes predicted not to form triple helices with any DEL_HCT_hs (Fig. 3F, Suppl. Table 1). This comparison was conducted under the conditions of an adjusted p-value < 0.01, Fold Enrichment ≥ 0.001, and |log2(Fold Change of Fold Enrichment)| ≥ 1. Consequently, it was revealed that ECM-related genes and cell growth-related genes were included among the DEL_HCT_hs target genes, whereas vesicle-related genes and genes associated with topologically incorrect proteins were included among the genes not targeted by any DEL_HCT_hs(Fig. 3H, Suppl. Table 2, 3). Among the top five GOBP terms for FoldEnrichment of DEL_HCT_hs target genes, the terms related to ECM contained genes that were not targeted by DEL_HCT_hs. However, the log2(Fold Change of FoldEnrichment) was 1.36, indicating an enrichment towards DEL_HCT_hs target genes. The remaining terms, which were significantly associated with cell growth and positive regulation of cell adhesion, did not include any genes among the non-targets of DEL_HCT_hs. Conversely, for the top five GOBP terms with the highest FoldEnrichment among non-target genes of DEL_HCT_hs, none of the significantly associated genes were found within the DEL_HCT_hs target genes.

## Discussion

Understanding the function of lncRNAs is crucial for a deeper comprehension of biological phenomena. LncRNAs have garnered significant attention in recent years due to their involvement in nearly all aspects of the central dogma, from the epigenome to protein stability(Herman et al., 2022). This involvement suggests their contribution to the evolution of life. However, previous research has primarily focused on their potential as biomarkers for diseases and drug targets(Nemeth et al., 2024), leaving their role in evolution largely unexplored. One notable difference between humans and closely related species, such as the Pan genus, is the presence of bipedalism. The relationship between lncRNAs and bipedalism remains unclear. Bipedalism is a result of structural changes in the lower limbs, with the skeleton being formed from cartilage during development. Given the skeletal differences in the lower limbs between humans and Pan, it is hypothesised that the cellular arrangement of cartilage in humans significantly diverges from that of other organisms during developmental stages. Although mature chondrocytes may not necessarily retain memories of their developmental phase, focusing on the distinctive RNA expression profiles of chondrocytes during development may elucidate genes involved in bipedal locomotion. Previously, we developed a method to differentiate iPS cells into chondrocytes by mimicking developmental stages(Yamada et al., 2021). However, this method is inherently artificial, making it challenging to discern whether the observed differences from mature chondrocytes are specific to the developmental stage or artifacts of the artificial process. Furthermore, the mechanisms of limb development are widely conserved, and the specific contributions of various genes at different stages are well-documented(Ornitz and Marie, 2015). In contrast, lncRNAs exhibit characteristics such as lower conservation compared to mRNAs, involvement in nearly all aspects of the central dogma, and in some cases, peptide coding. Therefore, by focusing on the functions of human-specific lncRNAs, it is possible to elucidate contributions to chondrocyte differentiation and, consequently, to bipedal locomotion from a perspective distinct from the conserved mechanisms of limb development. This study aims to investigate the expression of lncRNAs in cartilage during human development by comparing ExpLBM, as reported previously, with HCT differentiated from it, to infer the function of human-specific lncRNAs.

Initially, the functions of peptides encoded by DEL_HCT_hs were predicted, revealing that only 2 out of 15 peptides could be estimated, both related to responses to stimuli and stress (Suppl. Fig. 2). Furthermore, predictions of proteins that could bind to DEL_HCT_hs, followed by GO analysis of these proteins, indicated that they possess RNA splicing functions (Fig. 2C). Additionally, they were found to possess functions related to interactions with mRNAs. This study did not employ RNA-RNA interaction analyses such as hiCLIP or CLASH-Seq; thus, the specific genes targeted by human-specific lncRNAs for RNA splicing remain unknown. However, the highly ranked GO biological process terms such as RNA splicing, regulation of mRNA metabolic process, RNA transport, establishment of RNA localization, mRNA stabilization, and positive regulation of translation contribute to the structure and expression levels of proteins. Particularly, if human-specific splicing is regulated by DEL_HCT_hs, slight variations in protein function may be involved in bipedal locomotion. Investigating RNA splicing regulated by DEL_HCT_hs remains a task for future research.

Next, to investigate the potential regulatory role of DEL_HCT_hs on gene expression, the ability to form triple helices with DEG_HCT promoter regions was predicted (Fig. 3). The results demonstrated that promoters of genes related to cartilage formed triple helices (Fig. 3C, D), and that target genes of DEL_HCT_hs were enriched in ECM-related genes (Fig. 3H). The potential for triplex formation with human-specific genes among DEG_HCT was also investigated. It was demonstrated that triplexes could form with GSTT2B, ZFP36L1, and ANGPTL5, with a particular propensity for formation in the promoter region of ANGPTL5, which is predicted to be active in the extracellular matrix and extracellular space, including collagen. It is known that lncRNAs bind to nearby gene promoter regions to regulate expression(Dhaka et al., 2024), and among the lncRNAs predicted to form triple helices with DEG promoter regions, HSALNG0055279 and HSALNG0123261 were near SMOC2 and NFIC, respectively (Table 2). In mice, knockdown of SMOC1 along with SMOC2 results in impaired bone formation in the cranium, limbs, and mandible(Takahata et al., 2021). NFIC, a transcription factor, targets chondrocyte and joint marker genes. The mechanical stimuli transmitted by the ECM are relayed into the cell by various stimulus and stress response proteins, and it is intriguing that peptides derived from DEL_HCT_hs were predicted to respond to such stimuli and stress (Suppl. Fig. 2C). These findings suggest that human-specific lncRNAs may influence the composition of the ECM.

Another significant difference in skeletal structure between humans and other animals is the angle of the femur relative to the body axis. In chimpanzees and gorillas, close relatives of humans, this angle is 75°, whereas in humans it is 150°(Kozma et al., 2018), increasing the load on the joints. It has been revealed that Japanese macaques, known for their performance in “saru-mawashi” and trained to walk bipedally, adapt to the stresses of bipedal locomotion by increasing the strength of the trabecular bone in the femoral head and enhancing ligament attachments compared to their wild counterparts(Cazenave et al., 2024). Given that human-specific lncRNAs regulate the expression of ECM-related genes, it is suggested that these lncRNAs may have evolved to control ECM composition in response to the increased joint load due to bipedalism. This study focused only on the functionally estimated DEL_HCT_hs, but cellular and animal experiments on other lncRNAs might reveal those that have adapted to and contribute to bipedalism. Though this study primarily involved functional predictions using tools, empirical verification using cells and animals is necessary. Nevertheless, this research suggests that investigating lncRNAs from an evolutionary perspective could elucidate their role as drivers of cellular processes associated with evolution and adaptation.

## Conclusions

This study suggests that human-specific lncRNAs expressed during chondrogenesis may be associated with the ECM, potentially as an adaptation resulting from bipedalism. These findings have implications not only for evolutionary biology but also for therapeutic applications. Specifically, regulating human-specific lncRNAs could improve the quality of the ECM, potentially leading to the development of regenerative cartilage tissues that more closely resemble normal human joint cartilage. Furthermore, elucidating the mechanisms of evolution-related lncRNAs could contribute to understanding the pathogenesis of conditions such as osteoarthritis and neural tube defects, which are more prevalent in humans compared to other species.

## Supplementary Materials

Figure S1: Phylogenetic tree of biological classification

Figure S2: DEL_HCT_hs-Protein Interactions

Figure S3: Triplex Predictions of DEL_HCT_hs

Table S1: Fold enrichment for the GO analysis conducted on DEG_HCT, predicted to be targeted by DEL_HCT_hs in the promoter region, and other DEG_HCT.

Table S2: GO analysis results of DEG_HCT, predicted to be targeted by DEL_HCT_hs in the promoter region

Table S3: GO analysis results of DEG_HCT, predicted not to be targeted by DEL_HCT_hs in the promoter region

## Acknowledgments

Grants-in-Aid for Scientific Research from the Japan Society for the Promotion of Science (23K14384 to T. Osone; 23K08677 to T. Takao; 23K21368 to T. Takarada) and JST FOREST Program (JPMJFR225H to T. Takarada). These funders had no role in the study design, data collection and analysis, decision to publish, or preparation of the manuscript.

The computation was carried out using the General Projects on supercomputer “Flow” at Information Technology Center, Nagoya University.

## Conflicts of Interest

The authors declare no conflict of interest.

---

**Supplementary Figure 1.**
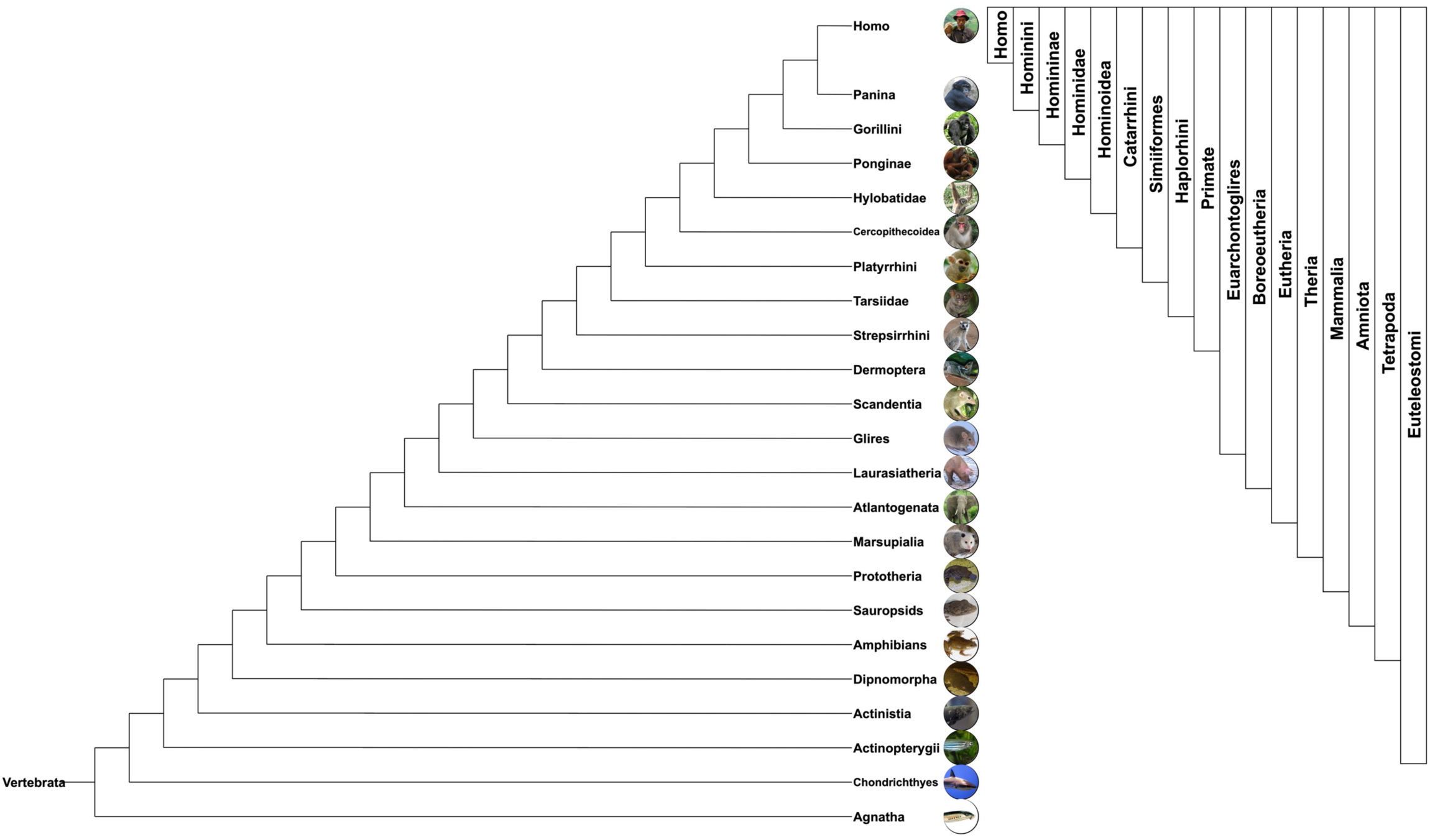
Phylogenetic tree of biological classification. The range indicated by the classification used in Fig. 1E is shown in a box.

**Supplementary Figure 2.**
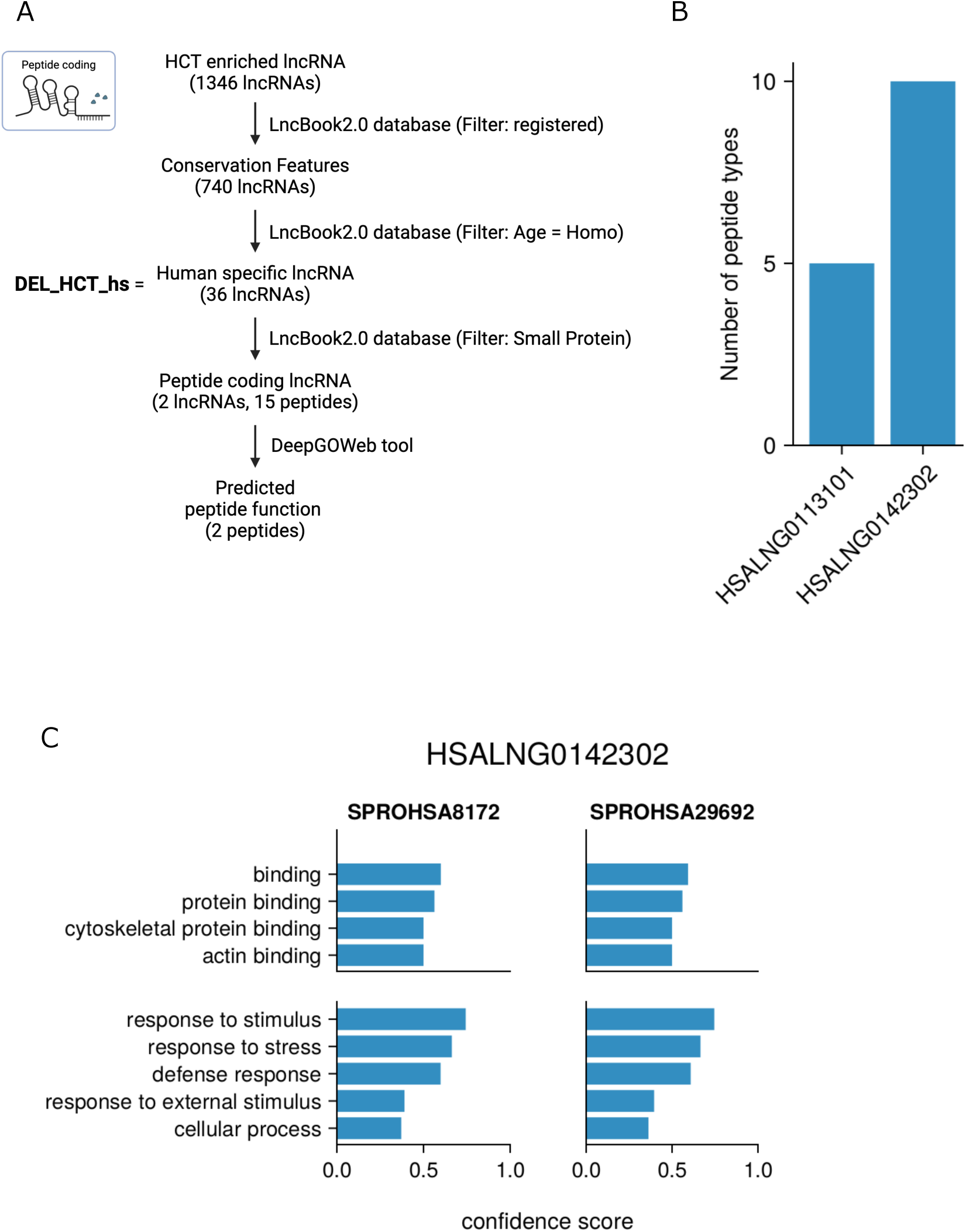
Predicted coding peptide functions of DEL_HCT_hs. (A) Schematic overview of the prediction methodology. (B) Bar graph showing lncRNAs reported to code for peptides and the number of peptides coded by these lncRNAs. (C) Bar graph of predicted peptide functions: the upper panel for Gene Ontology Molecular Function, and the lower panel for Gene Ontology Biological Process.

**Supplementary Figure 3.**
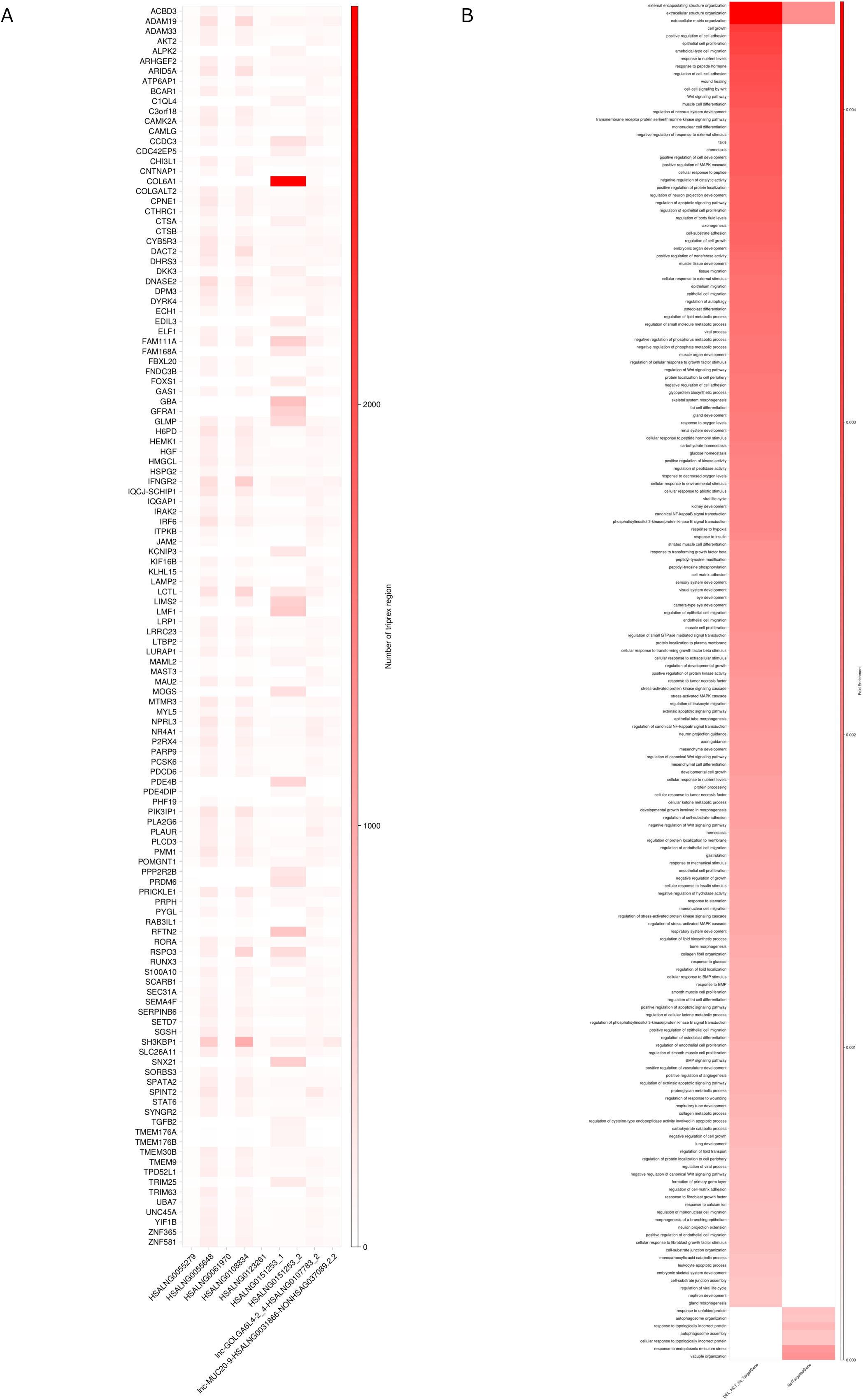
Triplex Predictions of DEL_HCT_hs. (A) Heatmap of formation sites for 9 DEL_HCT_hs predicted to form triplexes with DEG_HCT promoter regions. The top 50 formation sites for each lncRNA were extracted and combined into one heatmap. (B) Heatmap of formation sites for human-specific proteins, plotted using the raw values of region numbers instead of Z-scores.

## Notes

### Competing Interest Statement

The authors have declared no competing interest.

## References

Amatria-Barral, I., González-Domínguez, J., Touriño, J., 2023. PATO: genome-wide prediction of lncRNA–DNA triple helices. Bioinformatics 39, btad134. 10.1093/bioinformatics/btad134

Burrows, A.M., 2008. The facial expression musculature in primates and its evolutionary significance. BioEssays 30, 212–225. 10.1002/bies.20719

Cazenave, M., Nakatsukasa, M., Mazurier, A., M. Skinner, M., 2024. Identification of functionally related adaptations in the trabecular network of the proximal femur and tibia of a bipedally trained Japanese macaque. Anthropological Science 132, 13–26. 10.1537/ase.2307142

Dhaka, B., Zimmerli, M., Hanhart, D., Moser, M.B., Guillen-Ramirez, H., Mishra, S., Esposito, R., Polidori, T., Widmer, M., García-Pérez, R., Julio, M.K., Pervouchine, D., Melé, M., Chouvardas, P., Johnson, R., 2024. Functional identification of cis-regulatory long noncoding RNAs at controlled false discovery rates. Nucleic Acids Res 52, 2821–2835. 10.1093/nar/gkae075

Gil, N., Ulitsky, I., 2020. Regulation of gene expression by cis-acting long non-coding RNAs. Nat Rev Genet 21, 102–117. 10.1038/s41576-019-0184-5

Harcourt-Smith, W.H.E., 2010. The First Hominins and the Origins of Bipedalism. Evolution: Education and Outreach 3, 333–340. 10.1007/s12052-010-0257-6

Herman, A.B., Tsitsipatis, D., Gorospe, M., 2022. Integrated lncRNA function upon genomic and epigenomic regulation. Mol Cell 82, 2252–2266. 10.1016/j.molcel.2022.05.027

Hofman, M., 2014. Evolution of the human brain: when bigger is better. Front Neuroanat 8. 10.3389/fnana.2014.00015

Kamberov, Y.G., Guhan, S.M., DeMarchis, A., Jiang, J., Wright, S.S., Morgan, B.A., Sabeti, P.C., Tabin, C.J., Lieberman, D.E., 2018. Comparative evidence for the independent evolution of hair and sweat gland traits in primates. J Hum Evol 125, 99–105. 10.1016/j.jhevol.2018.10.008

Kozma, E.E., Webb, N.M., Harcourt-Smith, W.E.H., Raichlen, D.A., D’Août, K., Brown, M.H., Finestone, E.M., Ross, S.R., Aerts, P., Pontzer, H., 2018. Hip extensor mechanics and the evolution of walking and climbing capabilities in humans, apes, and fossil hominins. Proceedings of the National Academy of Sciences 115, 4134–4139. 10.1073/pnas.1715120115

Kulmanov, M., Zhapa-Camacho, F., Hoehndorf, R., 2021. DeepGOWeb: fast and accurate protein function prediction on the (Semantic) Web. Nucleic Acids Res 49, W140–W146. 10.1093/nar/gkab373

Kuo, C.-C., Hänzelmann, S., Sentürk Cetin, N., Frank, S., Zajzon, B., Derks, J.-P., Akhade, V.S., Ahuja, G., Kanduri, C., Grummt, I., Kurian, L., Costa, I.G., 2019. Detection of RNA–DNA binding sites in long noncoding RNAs. Nucleic Acids Res 47, e32–e32. 10.1093/nar/gkz037

Li, Z., Liu, L., Feng, C., Qin, Y., Xiao, J., Zhang, Z., Ma, L., 2023. LncBook 2.0: integrating human long non-coding RNAs with multi-omics annotations. Nucleic Acids Res 51, D186–D191. 10.1093/nar/gkac999

Long, F., Ornitz, D.M., 2013. Development of the Endochondral Skeleton. Cold Spring Harb Perspect Biol 5, a008334. 10.1101/cshperspect.a008334

Makałowski, W., Zhang, J., Boguski, M.S., 1996. Comparative analysis of 1196 orthologous mouse and human full-length mRNA and protein sequences. Genome Res 6, 846–857. 10.1101/gr.6.9.846

Nemeth, K., Bayraktar, R., Ferracin, M., Calin, G.A., 2024. Non-coding RNAs in disease: from mechanisms to therapeutics. Nat Rev Genet 25, 211–232. 10.1038/s41576-023-00662-1

Nishimura, T., Tokuda, I.T., Miyachi, S., Dunn, J.C., Herbst, C.T., Ishimura, K., Kaneko, A., Kinoshita, Y., Koda, H., Saers, J.P.P., Imai, H., Matsuda, T., Larsen, O.N., Jürgens, U., Hirabayashi, H., Kojima, S., Fitch, W.T., 2022. Evolutionary loss of complexity in human vocal anatomy as an adaptation for speech. Science (1979) 377, 760–763. 10.1126/science.abm1574

Noviello, T.M.R., Di Liddo, A., Ventola, G.M., Spagnuolo, A., D’Aniello, S., Ceccarelli, M., Cerulo, L., 2018. Detection of long non–coding RNA homology, a comparative study on alignment and alignment–free metrics. BMC Bioinformatics 19, 407. 10.1186/s12859-018-2441-6

O’Neill, M.C., Lee, L.-F., Demes, B., Thompson, N.E., Larson, S.G., Stern, J.T., Umberger, B.R., 2015. Three-dimensional kinematics of the pelvis and hind limbs in chimpanzee (Pan troglodytes) and human bipedal walking. J Hum Evol 86, 32–42. 10.1016/j.jhevol.2015.05.012

Ornitz, D.M., Marie, P.J., 2015. Fibroblast growth factor signaling in skeletal development and disease. Genes Dev 29, 1463–1486. 10.1101/gad.266551.115

Ramakrishnaiah, Y., Morris, A.P., Dhaliwal, J., Philip, M., Kuhlmann, L., Tyagi, S., 2023. Linc2function: A Comprehensive Pipeline and Webserver for Long Non-Coding RNA (lncRNA) Identification and Functional Predictions Using Deep Learning Approaches. Epigenomes 7. 10.3390/epigenomes7030022

Razmara, E., Bitaraf, A., Yousefi, H., Nguyen, T.H., Garshasbi, M., Cho, W.C., Babashah, S., 2019. Non-Coding RNAs in Cartilage Development: An Updated Review. Int J Mol Sci 20. 10.3390/ijms20184475

Richmond, B.G., Roach, N.T., Ostrofsky, K.R., 2016. Evolution of the Early Hominin Hand, in: Kivell, T.L., Lemelin, P., Richmond, B.G., Schmitt, D. (Eds.), The Evolution of the Primate Hand: Anatomical, Developmental, Functional, and Paleontological Evidence. Springer New York, New York, NY, pp. 515–543. 10.1007/978-1-4939-3646-5_18

RNAcentral: a comprehensive database of non-coding RNA sequences, 2017. . Nucleic Acids Res 45, D128–D134. 10.1093/nar/gkw1008

Statello, L., Guo, C.-J., Chen, L.-L., Huarte, M., 2021. Gene regulation by long non-coding RNAs and its biological functions. Nat Rev Mol Cell Biol 22, 96–118. 10.1038/s41580-020-00315-9

Takahata, Y., Hagino, H., Kimura, A., Urushizaki, M., Kobayashi, S., Wakamori, K., Fujiwara, C., Nakamura, E., Yu, K., Kiyonari, H., Bando, K., Murakami, T., Komori, T., Hata, K., Nishimura, R., 2021. Smoc1 and Smoc2 regulate bone formation as downstream molecules of Runx2. Commun Biol 4, 1199. 10.1038/s42003-021-02717-7

Wang, J., Zhao, Y., Gong, W., Liu, Y., Wang, M., Huang, X., Tan, J., 2021. EDLMFC: an ensemble deep learning framework with multi-scale features combination for ncRNA–protein interaction prediction. BMC Bioinformatics 22, 133. 10.1186/s12859-021-04069-9

Yamada, D., Nakamura, M., Takao, T., Takihira, S., Yoshida, A., Kawai, S., Miura, A., Ming, L., Yoshitomi, H., Gozu, M., Okamoto, K., Hojo, H., Kusaka, N., Iwai, R., Nakata, E., Ozaki, T., Toguchida, J., Takarada, T., 2021. Induction and expansion of human PRRX1+ limb-bud-like mesenchymal cells from pluripotent stem cells. Nat Biomed Eng 5, 926–940. 10.1038/s41551-021-00778-x

Zhu, J., Yu, W., Wang, Y., Xia, K., Huang, Y., Xu, A., Chen, Q., Liu, B., Tao, H., Li, F., Liang, C., 2019. lncRNAs: function and mechanism in cartilage development, degeneration, and regeneration. Stem Cell Res Ther 10, 344. 10.1186/s13287-019-1458-8

